# Analysis of multicellular anatomical structures from spatial omics data using sosta

**DOI:** 10.1101/2025.10.13.682065

**Authors:** Samuel Gunz, Helena L. Crowell, Mark D. Robinson

## Abstract

Spatial omics technologies enable high-resolution, large-scale quantification of molecular features while preserving the spatial context within tissues. Existing analysis methods largely focus on spatial arrangements of single cells, whereas biological function often emerges from multicellular arrangements. Here, we introduce structure-based analysis of spatial omics data, which focuses on the direct analysis of multicellular, anatomical structures. We illustrate this type of analysis using two publicly available datasets and provide `sosta`, an open-source Bioconductor package for broad community use.

## Introduction

Spatial organisation reflects biological function across scales. Cells and tissues tend to self-organize into spatially segregated anatomical structures with distinct biological function, which is often reflected in the morphology and location of such multicellular structures and organs [1] (Fig 1a-b). Recent advances have enabled researchers to quantify molecular features at an unprecedented scale and resolution, while preserving their spatial context [2] (Fig 1c). This motivates a data analysis approach that focuses on the anatomical structure dimension of spatial omics data.

**Fig 1.**
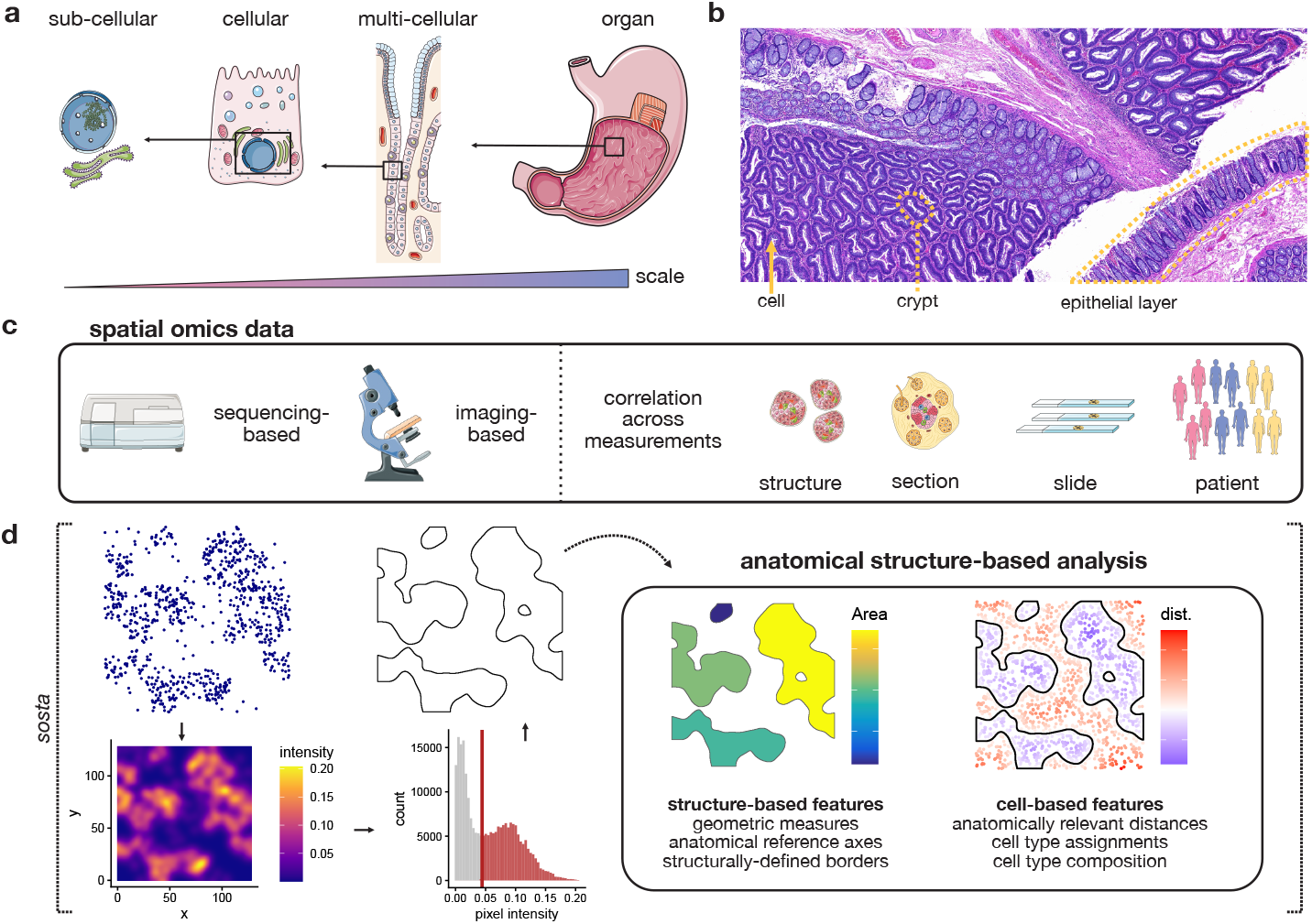
Structure-based analysis of spatial omics data. **a**) Spatial arrangements reflect functions across scale, ranging from the subcellular to the organ level. **b**) An example histopathology image that illustrates diverse multicellular anatomical structures in the transition from healthy intestinal epithelium to premalignant lesions (taken from [10]). **c**) Spatial omics technologies are used to study biological phenomena across different scales, ranging from subcellular to multicellular, with correlation also present at different scales. **d**) Illustrated workflow of density-based reconstruction of anatomical structures using a threshold on the point pattern intensity of cell centroids as implemented in the `sosta` package, followed by the analysis of anatomical structures by quantifying geometric features, or by using anatomical structure information as a reference scale for downstream analysis. Illustrations in **a** and **c** were adapted from [Servier Medical Art](https://smart.servier.com/), licensed under [CC-BY 4.0](https://creativecommons.org/licenses/by/4.0/).

While many computational methods have been proposed to analyse emerging types of spatial omics data [3], they usually focus on the arrangement of individual cells, often within a single tissue or slide; for example, the analysis of cell neighbourhoods, spatial domains, or cell-cell interaction patterns. Moreover, methods to detect spatially variable features or gradients often do not account for heterogeneity at the level of anatomical structures [2, 4]. In addition, few methods exist for the analysis of multi-sample multi-condition spatial datasets [3]. To date, general frameworks and strategies are lacking to perform analysis at the (anatomical) structure level, where multicellular arrangements become the focus of analysis. Some early attempts have been made, such as extracting spatial domains and quantifying geometric features [5, 6]. Here, we introduce a new genre of data analysis, so-called structure-based analysis, whereby (computationally-extracted) anatomical structures represent the basic unit and resolution to analyse the data at (Fig 1c-d). As such, we complement ongoing research in cell type annotation [7] and spatial domain detection (spatial clustering) [8] since the generated cell type or domain labels are possible inputs for the reconstruction of spatial structures.

We demonstrate this type of analysis using two datasets: quantifying structural rearrangements during colorectal malignancy transformation; and, recovery of structurally relevant gene expression gradients in human tonsil germinal centres. To build a foundation for such analyses, we also introduce `sosta` (<u>s</u>patial <u>o</u>mics <u>st</u>ructure <u>a</u>nalysis), an open-source Bioconductor [9] package for reconstruction, characterization, and comparison of anatomical structures from spatial omics data across scales.

## Design and Implementation

Using a structure-based analysis approach, we aim to bridge the gap between classical histopathology and spatial omics by directly quantifying structural rearrangement and subsequent changes in the molecular landscape. To characterize structural differences, we rely on a structure reconstruction implemented in our package `sosta`. This approach allows for flexible and fast “segmentation” of a set of (disconnected) anatomical structures by relying on coordinates of associated features of interest, such as cell types or transcript assignments (Fig 1b). We then focus on structure-based analysis by quantifying a set of metrics related to the recovered anatomical structures and using this resolution to examine the data (Fig 1d).

### Density-based reconstruction of anatomical structures

In `sosta`, reconstruction of the outlines of anatomical structures is performed by using a cut-off on a point pattern density estimate, where the point pattern corresponds to either the centroids of selected cell type(s), the locations of transcripts of interest, features marking a spatial domain (or cluster or functional pathway), or any other spatial covariate associated with individual cells. As previously described by Diggle et al. [11], a density image of a point pattern can be estimated by

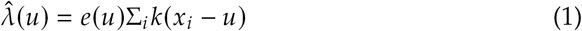

where 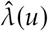 is the estimated intensity at point *u, k*(·) is a Gaussian kernel and *e* ( *u* ) is an edge correction factor.

By default, the bandwidth of the smoothing kernel is estimated using cross validation on the bandwidth selection as described by Diggle et al. [11]. The density estimation and kernel selection are implemented in the R package `spatstat` [12]. To facilitate the use of this approach, we implemented our package within the R/Bioconductor environment and work with common data structures for spatial (omics) data. The resulting anatomical structures are stored as polygons using the `sf` package [13].

### Parameter estimation

To estimate the cut-off threshold value *c* for reconstruction, we use an automated estimation of the threshold 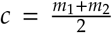 where *m*_1_ and *m*_2_ are the density values at the first and second mode of the intensity distribution over all pixels in the estimated density image. In settings where the distribution is unimodal, the cut-off value lies at the peak of the distribution. We further provide auxiliary functions to facilitate manual adjustment and optimization of this cut-off value, as well as the bandwidth of the Gaussian kernel *k*. In particular, we provide functions to visualise density images, the intensity distributions, and to estimate the range and variability of automatically estimated parameters. Overall, we noticed that downstream results were consistent across a range of parameter settings (i.e., bandwidth of smoothing kernel and cut-off threshold), indicating robustness of our reconstruction approach. However, at an upper or lower range of values we observed substantial deviations, especially in the number of detected structures (Fig S3c).

### Quantification of morphological features

After the anatomical structures have been reconstructed, `sosta` offers functions to calculate a range of geometric features such as the area, circularity or eccentricity of each individual structure. An overview of all structural metrics calculated by `sosta` given in Suppl. Table S1.

### Cell-based features

In addition to morphological features, `sosta` offers functions that relate individual cells to anatomical structures. They compute intersections between cells and structures and measure distances to structure borders. The functions support quantifying cell type composition dynamics and defining gradients relative to structure border or anatomical landmarks.

### Differential feature analysis

Multi-sample multi-condition analysis of spatial omics datasets requires care as this type of data often harbours multiple levels of correlation. In the case of structure-based analysis, we can have multiple relevant experimental units such as multiple structures in a section, multiple sections in one slide, multiple slides per patient and multiple patients per condition. In downstream analysis, it is critical to recognize that these are not independent experimental units. For differential feature analysis, linear mixed effects modelling tools offer effective strategies to account for nested correlation structure [14]. An analysis of a multi-sample multi-condition spatial proteomics dataset by Damond et al. [15] revealed strong correlation effects within experimental units at different levels. Furthermore, using a semi-synthetic simulation framework adapted from Yu et al. [14] we noticed that failing to account for the nested correlation structure in the Damond et al. dataset lead to an increase of the false discovery rate (FDR) of up to 20% while mixed effect models retained the FDR below the selected 5% level (Suppl. Fig S1, Suppl. Methods).

## Results

### Methods comparison

Other methods exist that take coordinates and labels as input and output a polygon that can be used for downstream morphological quantification and subsequent geometric operations. We compared `sosta`against the R/Bioconductor packages `imcRtools` [16] and `SPIAT` [17], and the Python package `CellCharter` [5] that use a concave hull around a defined set of points to define structures and to the Python package `GRIDGENE` [6] that employs a point pattern density-based approach conceptually similar to ours (Suppl. Methods). We adapted the code of `SPIAT` as it did not include any function that outputs polygons but only calculates them internally. Among these methods, `GRIDGENE` does not include automatic estimation of parameters for structure reconstruction so that all parameters require manual adjustment.

Since ground truth datasets with expert annotations often exhibit subjective annotation bias [8], we based our comparison on simulations. We simulated binary images using random seeds and a Gaussian smoothing kernel to define structures and subsequently sampled points within these structures using a Poisson process with varying intensity, including background noise with varying strength. We then performed reconstruction using the different methods and converted the resulting polygons into binary images. Method performance was evaluated using the Jaccard index between the original binary image and the reconstruction (Suppl. Fig S2, Suppl. Methods).

Density based methods clearly outperformed approaches that construct a concave hull around a set of points. `sosta` achieved the best performance, with `GRIDGENE` reaching similar results. However, this occurred only when selecting the best `GRIDGENE` result across a range of parameters, using the highest Jaccard index against the ground truth. The performance of `sosta` showed a substantially higher variance in high noise settings compared to low noise settings, but the average performance was comparable in both settings. The number of target cells had little influence on the reconstruction accuracy. Overall, these results highlight the advantage of density-based methods and automated parameters selection as implemented in `sosta` (Suppl. Fig S2c-e).

### Reconstruction and quantification of epithelial crypts in colon tissue

To highlight structure-based analysis of spatial transcriptomics data, we first quantified structural changes in the transformation from healthy human colon (HC) tissue to colorectal cancer (CRC) [10] (Fig 2a, Suppl. Fig S3a-c). This transition from HC crypt epithelial cells to CRC is a process that includes intermediate stages, such as polyps or tubulovillous adenomas (TVAs). These stages are typically identified by histopathology since they are characterized by discernible cellular morphology and structural rearrangement. The transcriptional dynamics along this transition have been studied in Crowell et al. [10].

**Fig 2.**
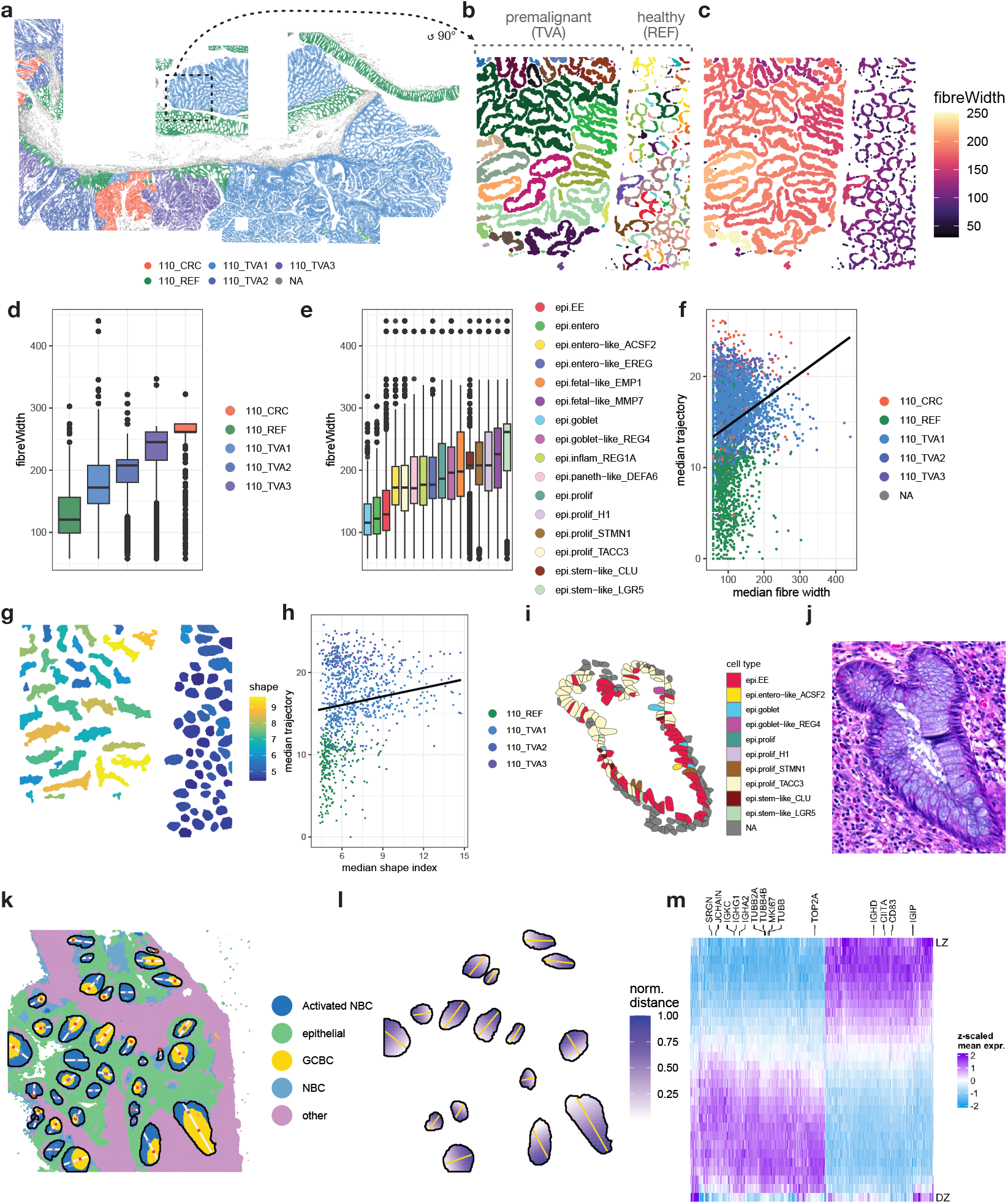
Two examples of structure-based analysis of spatial transcriptomics data. **a**) Overview of section 110 from Crowell et al. [10] (REF = reference / healthy colon, TVA = TVA regions, CRC = colorectal cancer regions). **b**) Subregion indicated by the dotted box in **a**, rotated counter-clockwise by 90°, that reflects the transition between normal colonic crypts and premalignant TVA regions. Results of the `sosta` density-based reconstruction are randomly coloured to highlight unique structures. **c**) Epithelial structures in b) coloured by their respective fibre width. **d**) Boxplot of the distribution of fibre width per cell (each cell was assigned the fibre width metric of its corresponding epithelial structure) grouped by the assigned region. **e**) Boxplot of per-cell fibre widths grouped by cell types. x-axis ordered by median fibre width. **f**) Association between the median fibre width per epithelial structure and its median (inferred) pseudo-time [10]. **g**) Reconstructed epithelial crypts of b). The colour reflects their corresponding “shape index”. **h**) Association between the median pseudo-time and the median shape index per reconstructed epithelial crypt. **i**) Examples of a *de novo* identified transition crypts using structure-based analysis and **j**) corresponding histopathology image (adjacent section). **k**) Structure-based analysis of germinal centres (GCs) in the human tonsil [20]; black outlines correspond to reconstructed GCs, colours reflect inferred clusters. Coloured squares correspond to reference points used to reconstruct anatomical reference axes (white lines). **l**) Reference axes through GCs; colours correspond to distances scaled between 0 and 1 within each GC. Heatmap of 1569 genes with spatially-varying expression within the GCs. Highlighted are known marker genes. LZ = light zone, DZ = dark zone.

To accentuate morphological changes between HC and (pre-)malignant regions, we calculated a range of geometric features using `sosta` and centred on a geometric metric called “fibre width”, which represents the width of the between inner and outer crypt border (Fig 1c, Suppl. Table S1, Methods). Fibre width increased along the pathological progression from HC to TVA to CRC, both at the structure and cell type level (Fig 2d-e). As such, this metric reflects the gradual morphological changes related to increase in crypt thickness. In healthy tissue, epithelial cells are lining the crypt; in hypertrophy and hyperplasia, there is an increase in size and number of epithelial cells, respectively. Next, dysplasia denotes a known premalignant state in which epithelial cells lose differentiation, present abnormal size and shape and lack polarization followed by carcinoma [18, 19]. To further study this progression, we compared the association of fibre width with the median pseudo-time per structure as inferred in the original publication for each cell. We observed a strong positive correlation between the median fibre width per epithelial structure and the median pseudo-time of cells within the structure (Spearmans’s *ρ* = 0.46, *p*-value < 10^−10^, Fig 2f). In addition to structural analysis of the epithelial cell layer, we further reconstructed and quantified individual epithelial crypts (Fig 2g). We found that the “shape index”, a morphological measure of a shape’s regularity, reflects the transition from regular crypts in the normal epithelium to irregular ones in premalignant regions (Spearmans’s *ρ* = 0.22, *p*-value < 10^−10^, Fig 2h). Furthermore, we used the reconstructed crypts to identify local transitions between normal and premalignant tissue, i.e., transition crypts. By filtering based on the frequency of cell types adjacent to the reconstructed crypts, we recovered one of seven manually annotated transition crypts and identified potentially *de novo* transition crypts. Although this approach failed to detect several manually annotated crypts, it revealed additional candidate crypts that also showed transition patterns in the histopathology image (Suppl. Methods, Fig 2i-j, Suppl. Fig S3d-h).

By using structure-based analysis, we gained additional insight about the morphological transition during the progression to CRC. For illustration, this was done on a single slide. However, this type of analysis could also be performed with multiple replicates and across conditions. In this case, it is crucial to account for correlation of the repeated measurements (i.e., structures) on the same experimental unit (i.e., sample) (Fig 1c).

### Reconstruction and analysis of anatomically relevant gradients in germinal centres of the human tonsil

In addition to geometric features, anatomical structures offer a baseline for structure-aware analysis of gene expression. Unlike conventional spatially variable gene analysis, this approach relates spatial variation of gene expression to meaningful anatomical axes. To demonstrate this, we performed structure-based analysis of a spatial transcriptomics dataset of human tonsil tissue [20] (Fig 2k, Suppl. Fig S4). Relevant anatomical structures in this dataset are germinal centres (GCs), locations where B cells mature along a defined anatomical trajectory: from light to dark zone (L/DZ) [21]. We used our density-based reconstruction approach to define GCs and, within each GC, constructed anatomically relevant axes by first identifying anatomical zonations in each GC then projecting a matching axis through each GC (Fig 2l, Suppl. Fig S4, Suppl. Methods). This allowed for quantification of gene expression along a unified anatomically relevant gradient across GCs.

Next, we averaged expression of all GC spots within binned distances and used the maximal information coefficient (MIC) [22], a metric that can capture linear and non-linear relationships, to identify genes that showed gene expression changes along the LZ-DZ axis. We identified a total of 1569 genes that showed a spatially varying gene expression profile (MIC > 0.999). These included known markers of B cells proliferation, maturation, and antigen presentation (e.g., MKI67, CIITA) [21, 23] (Fig 2m).

In this example, structure-based analysis offered a straightforward approach to recover interpretable and anatomically relevant gene expression gradients without relying on discrete cell types.

## Availability and Future Directions

To date, spatial omics analysis methods have primarily focused on the single-cell level, often failing to fully capture heterogeneity arising from differences in tissue architecture and cellular composition. Previous work has shown that scale is an important factor when analysing spatial omics data [24, 25]. This motivates a structure-based analysis approach that is aware of multicellular anatomical patterns, as has been shown in classic histopathology for decades [26]. Our package `sosta` functions as a modular and extensible tool within the Bioconductor framework to perform structure-based analysis at different scales. We provide comprehensive vignettes to enable a user friendly experience at https://bioconductor.org/packages/sosta.

In addition, our density-based reconstruction approach allows for analysis of anatomical structures ranging from few cells or transcripts such as tertiary lymphoid structures, up to the analysis of cancer tissue comprising thousands of cells. In these cases, the structure of interest were known in advance, but the approach we use and facilitate is a general one. Moreover, the type of structure-based analyses we have shown are applicable to a diverse set of biological tissues and questions. To date, researchers often use prior biological knowledge to guide selection of metrics that quantify relevant aspects in their data. This offers further opportunities to investigate unsupervised methods to first identify and quantify biologically relevant features at the anatomical structure level [27]. By leveraging existing statistical frameworks for analysis of structure-derived features, we show how structure-based analysis can ease biological discovery. However, researchers must heed the correlation structure of structure-based analysis of spatial omics data, since multiple measurements from the same experimental unit are extracted.

While our framework offers methods to reconstruct, quantify and compare anatomical structures ranging from single to multi-sample multi-condition setups, we acknowledge that there exists a plethora of methods that can be applied in each step of structure-based analysis. First, suitable targets for reconstruction have to be identified. These could range from marker expression, cell type labels, pathway signatures to spatial domains. Second, a suitable reconstruction method and parameters have to be chosen. Third, the range of metrics for downstream analyses have to be identified. For example, a range of methods exists to reconstruct and segment (multi-)cellular structures, the space of possible metrics that quantify anatomical structures is vast and different strategies to detect gradients exist [28]. Nonetheless, our flexible and extensible framework can accommodate future developments in this space.

## Code and data availability

`sosta` is freely available from Bioconductor since the 3.21 release https://bioconductor.org/packages/sosta. Code to reproduce analyses in this work are available at https://github.com/sgunz/sosta-manuscript.

The CosMx TVA dataset is available at https://zenodo.org/records/15574384 [29]. The IMC pancreatic islets dataset is available at https://data.mendeley.com/datasets/cydmwsfztj/2 and through Bioconductor’s ExperimentHub via the R `package imcdatasets` [15]. The Visium HD human tonsil (fresh frozen) dataset is available from the 10x Genomics website (https://www.10xgenomics.com/datasets) [20].

## Acknowledgments

We thank all members of the Robinson lab for helpful comments and feedback. We thank Martin Emons for continuous feedback and support, Izaskun Mallona and Giulia Moro for helpful insights into anatomy and biology, Sanne Verheul for feedback on the software, and Bianca Dumitrascu and Tammy Qiu for suggestions on the analysis.

Medical illustrations in Fig 1 were adapted from [Servier Medical Art](https://smart.servier.com/), licensed under [CC-BY 4.0](https://creativecommons.org/licenses/by/4.0/).

## Artificial Intelligence Tools

We acknowledge the use of following generative artificial intelligence tools in this work. *ChatGPT* (OpenAI, version 5.2 and lower) was used to support literature search, review individual paragraphs, suggest alternative phrasing for individual sentences, assist in generating and restructuring code for the `sosta` package and analyses, draft code documentation and help write unit tests. *DeepL Write* (free mode) was used to suggest alternative phrasing for individual sentences. All code was manually checked and we declare full responsibility for all content in this work.

## Author Contributions

**SG:** Conceptualisation, Methodology, Analysis, Visualisation, Software, Writing - Original Draft; **HLC:** Conceptualisation, Methodology, Writing - Original Draft, Supervision; **MDR:** Conceptualisation, Methodology, Writing - Original Draft, Supervision, Funding acquisition.

## Supporting Information

**S1 File Supplementary Information** Combined Supplementary Information including Supplementary Figures, Tables, Software Availability and Installation, and Supplementary Methods.

