## Supplementary Information for "Analysis of multicellular anatomical structures from spatial omics data using sosta"

#### Contents

**Supplementary Figures** 1

**Supplementary Table** 6

**Software Availability and Installation** 7

**Supplementary Methods** 7

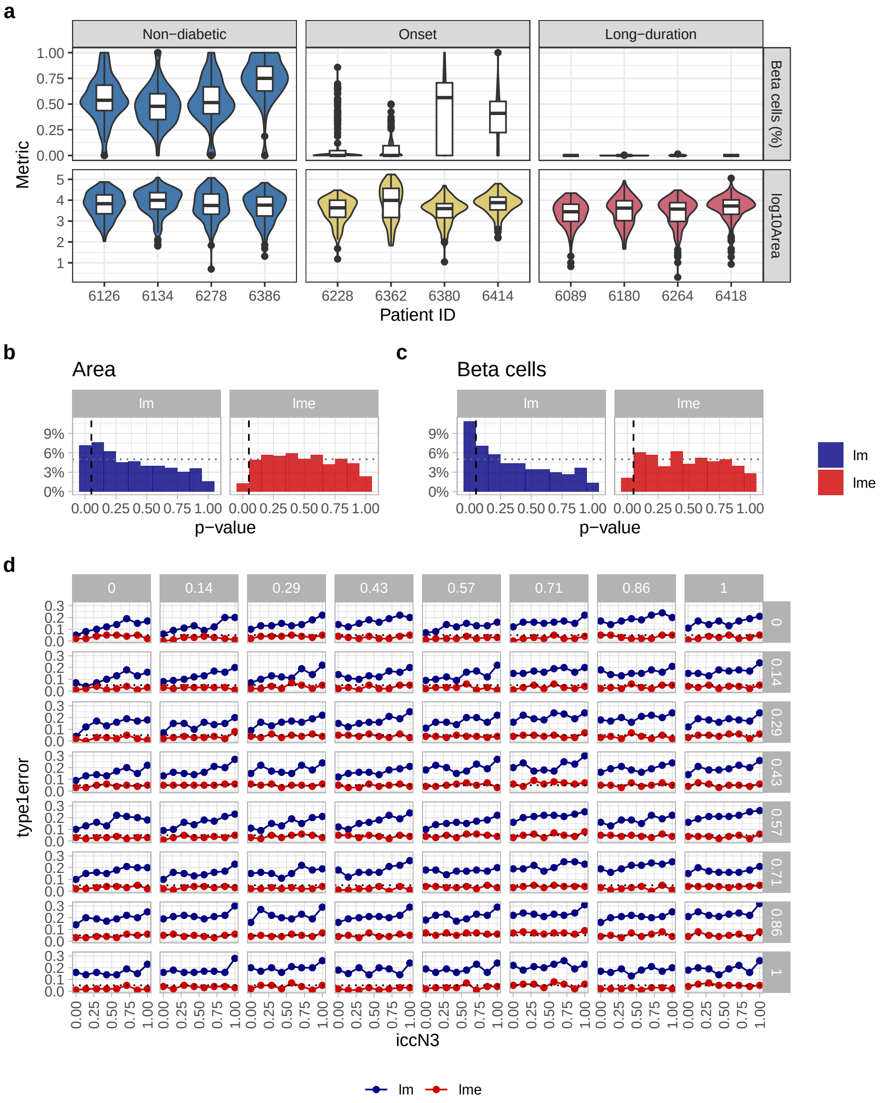

**Fig S1. Multi-sample multi-condition structure based analysis.** **a**) Distribution of three different metrics related to spatial structures across patients and conditions in an IMC dataset of pancreatic islets [5]. **b, c**) Histogram of p-values of repeated null simulations (no differences between conditions) based on intra class correlation coefficients estimated from the [5] dataset. Analysis shown for area of islets (b) and fraction of beta cells within islets (c). Dashed and dotted line indicate 5% threshold. **d**) Relationship between ICC for three different groups / conditions (x, y facets and x axis) and type 1 error for linear and linear mixed effect models. lm, linear model; lme, linear mixed effects model.

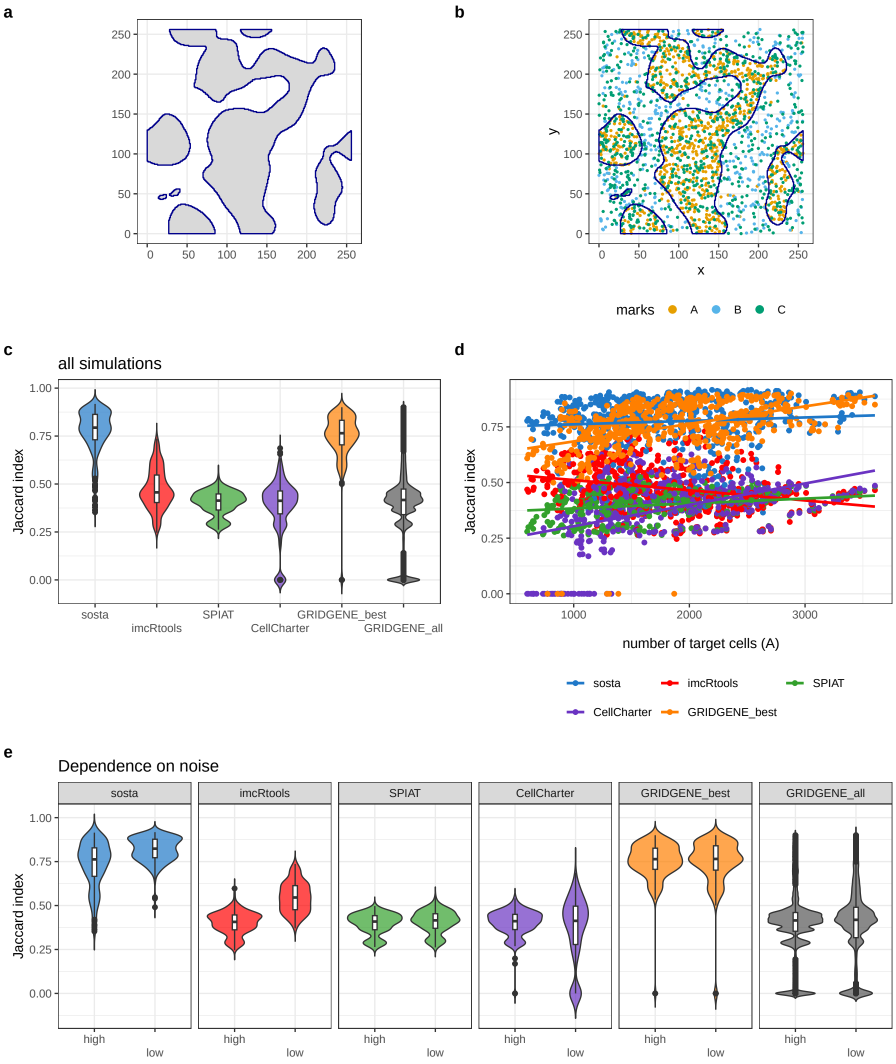

**Fig S2. Methods comparison. a**) Simulation of a binary image that mimics multi-cellular anatomical structures using the function `simulateTissueBlobs` of `sosta`. Gray regions outlined in blue represent tissue regions. **b**) Based on the binary image points that represent cells in the tissue are sampled (in-tissue) and background are sampled based on a Poisson process with different intensities. Mark "A" represents the tissue and was selected for reconstruction. **c**) Distribution of Jaccard indices (JIs) for all simulations and the different methods. `GRIDGENE` was split into the best parameter combination and the distribution of all parameters. **d**) Relationship between the number of target cells and the JI. **e**) Distribution of JIs for all simulations and the different methods split into low noise `noise_A < 0.01` and high noise `noise_A ≥ 0.01` scenarios.

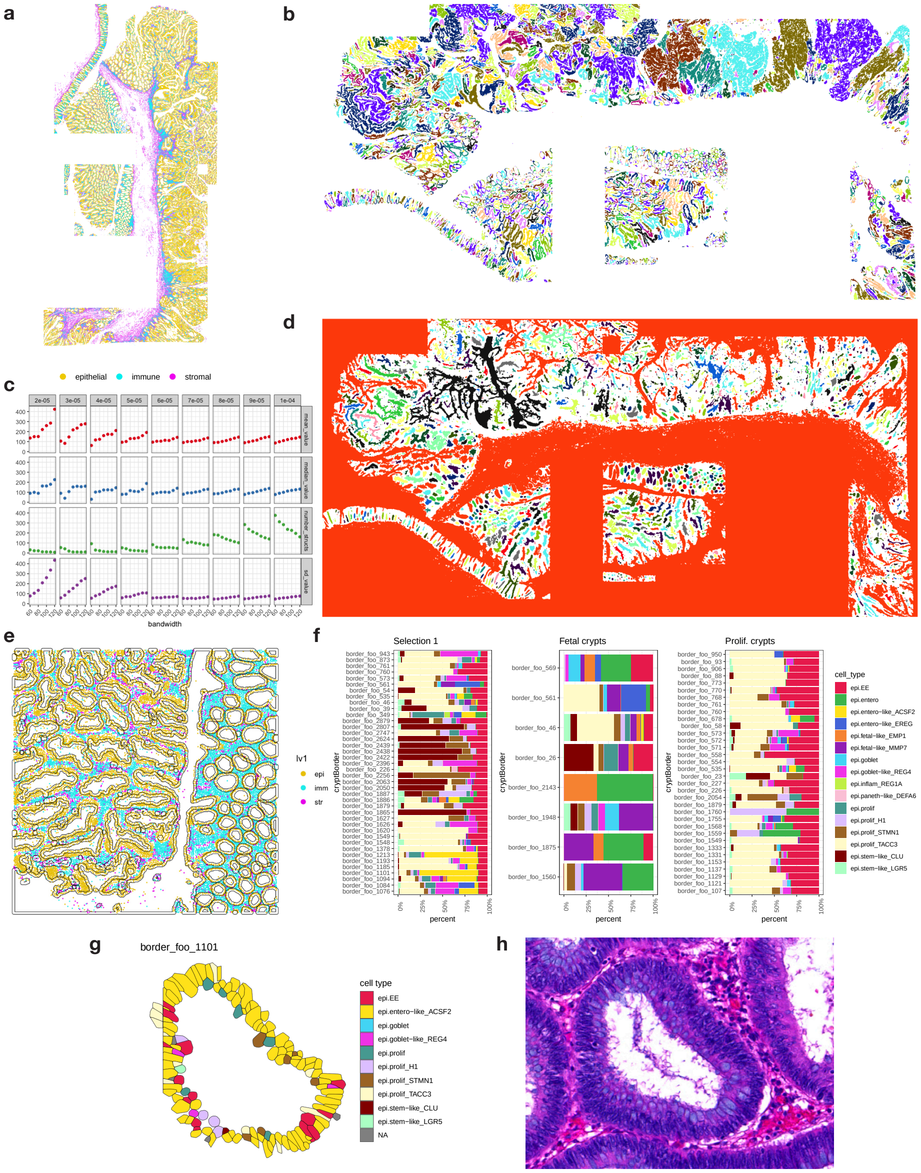

**Fig S3. Structure-based analysis of epithelial structures in the [1] dataset. a**) Slide 110 of the Crowell et al. [1] dataset. Colours correspond to cell type annotations of the original publication. **b**) All reconstructed epithelial structures using the density-based reconstruction method of `sosta`. **c**) Impact of parameters bandwidth (x) and threshold (facets) in reconstruction using `reconstructShapeDensitySPE` on the the mean, median and standard deviation of fibre width, and the number of recovered structures in a representative region (c.f. Fig. 2b). **d**) All reconstructed inter-epithelial (crypt) regions before filtering. **e**) Expansion of crypts to assign cells to crypts before selection of transition crypts. A representative region was chosen for illustration. **f**) Cell type composition of identified transition crypts. **g** Region in the histopathological image corresponding to the identified transition crypt in Fig 2i). **h**) Misidentified transitions crypt (false positive) and **g**) corresponding region in the histopathological image. CRC = colorectal cancer, REF = reference (healthy tissue), TVA = tubulovillous adenoma.

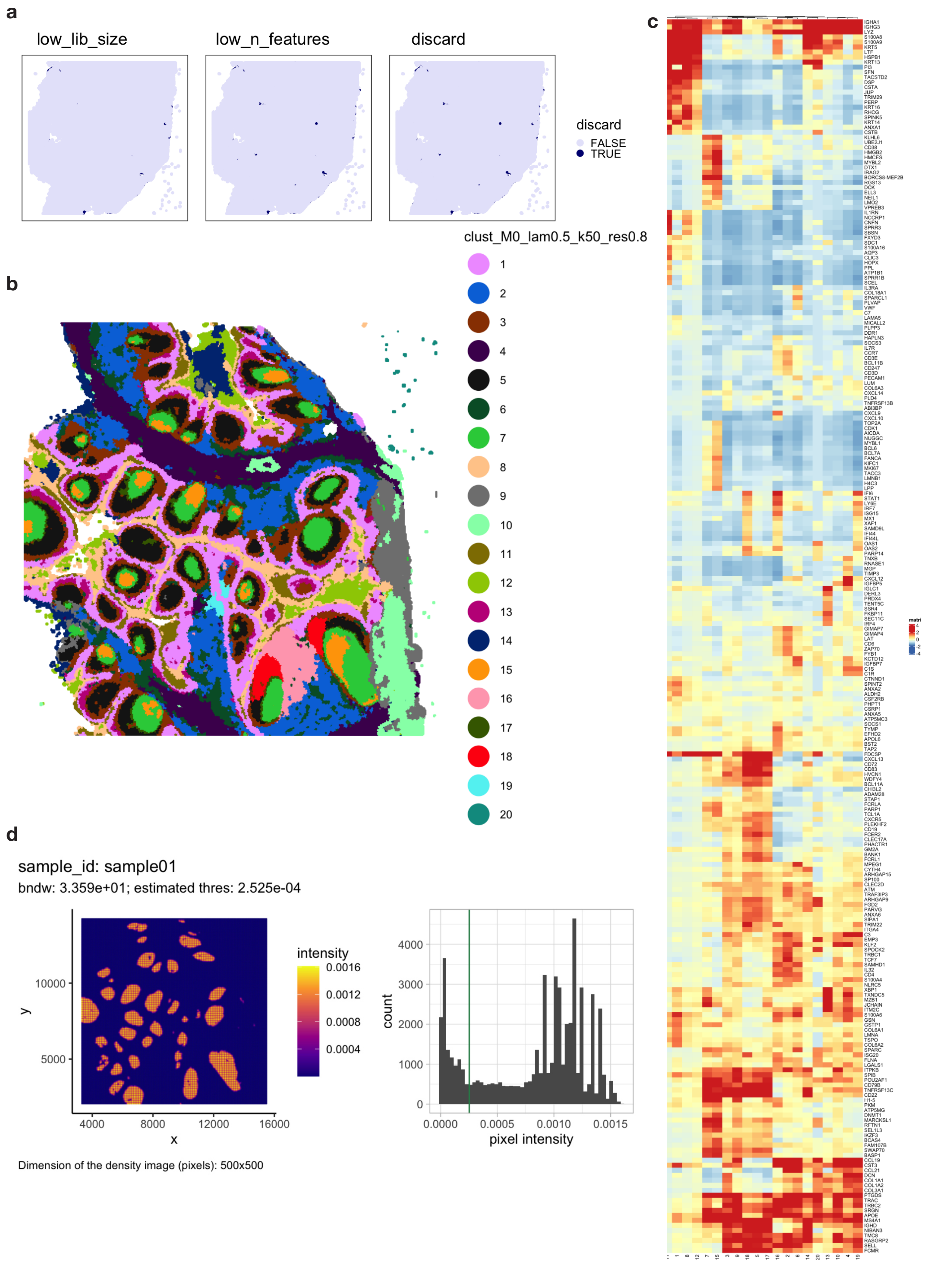

**Fig S4. Structure-based analysis of germinal centres in the human tonsil a)** Results of quality control using `SpotSweeper` [2] and different quality control (left to right: library size below threshold, number of features below threshold, combined flagged spots to discard). **b**) Results of clustering using `Banksy` [3]. **c**) Identified marker genes for the *Banksy* clusters using `findMarkers` of `scater` [4]. **d**) Density image showing density of GC cells with reconstruction parameters estimated by `sosta`.

#### Supplementary Table

| Metric | Formula | Function / Meaning |
|---|---|---|
| Area ($A$) | $A$ | |
| Perimeter ($P$) | $P$ | Length of the structure boundary. |
| Compactness ($C$) | $C = \frac{4\pi A}{P^2}$ | Normalized measure of compactness (1 = perfect circle). |
| Major Axis ($a$) | $a$ | Length of the long axis of the minimum bounding rectangle. |
| Minor Axis ($b$) | $b$ | Length of the short axis of the minimum bounding rectangle. |
| Eccentricity ($E$) | $E = \frac{b}{a}$ | Ratio of minor to major axis. 1 = circular, 0 = elongated. |
| Circularity / Roundness ($R$) | $R = \frac{4\pi A}{P_c^2}$ | Measures roundness using the convex hull perimeter $P_c$. Higher = more circular. |
| Solidity ($S$) | $S = \frac{A}{A_c}$ | Ratio of polygon area to convex hull area $A_c$. Measures convexity/density (1 = convex). |
| Curl | $\text{Curl} = 1 - \frac{a}{L_f},$ | Shape bending/curvature measure. $L_f$ is the effective fibre length along the main axis. |
| Fibre Length ($L_f$) | $L_f = \frac{4A}{P - \sqrt{P^2 - 16A}}$ | Characteristic length of the shape along the main axis. |
| Fibre Width ($W_f$) | $W_f = \frac{A}{L_f}$ | Characteristic width of the shape perpendicular to the main axis. |

**Table S1. List of geometric features calculated by *sosta*.**

### Software Availability and Installation

`sosta` is available through Bioconductor (v3.21 and higher). Details on the installation can be found in the corresponding vignettes `https://bioconductor.org/packages/release/sosta`.

Analyses in this manuscript were conducted using R version 4.5.1 and Bioconductor version 3.22 [6]. Results were visualised using `ggplot2` [7]. Packages for analyses in this manuscript were managed using `renv`. Details on the installation of R and Python can be found at `https://github.com/sgunz/sosta-manuscript`.

### Supplementary Methods

#### Multi-sample multi-condition structure-based analysis

Structure-based analysis in multi-sample multi-condition data adds a layer of complexity, as we will illustrate in an analysis of a spatial proteomics imaging mass cytometry (IMC) dataset of [5]. This dataset characterizes pancreatic islets in the course of type-one diabetic progression across twelve patients and three conditions. As such, it serves as a good example for structure-based analysis since it contains well characterized anatomical structures, i.e., the islets, where their biology is well studied.

We first use our point-pattern density reconstruction approach implemented in `sosta` to reconstruct pancreatic islets and quantify corresponding biologically relevant features, ranging from geometric measures, such as the area of the islets to biologically relevant metrics such as the proportion of different cell types. To identify biologically relevant features, we then perform feature-wise differential analysis on the structure derived features.

**Challenges of differential analysis** Notably, two aspects of this dataset are characteristic of the challenges when performing differential analysis of structure-derived features. First, the different scale of different metrics. Structure-derived metrics can be positive-continuos (e.g., area), bound between zero and one (e.g., eccentricity) or discrete (e.g., number of observations within a structure). Note that some features can be represented differently, which impacts downstream modelling (e.g., cell types within a structures can be expressed both as fractions, i.e., bounded between 0 and 1, or using positive discrete values representing counts of cells). Second, one should be mindful of the correlation structure. Since the dataset contains multiple islets per slide, multiple slides per patient and multiple patients per condition, there are indeed multiple levels of correlation structure. Therefore, features derived from individual islet structures are not independent measurements (Fig S3a). To quantify the correlation structure, we used the interclass correlation coefficient to measure correlation at different levels and generated simulations to assess the impacts of standard differential analysis. Not accounting for such correlation structures can be problematic and can lead to an inflated type I error (false positive) rate. For more discussion on this problem and an introduction to both linear and linear mixed-effect models, we refer the reader to [19].

**Estimate Correlations using ICC** To estimate correlation at different levels, we use the interclass correlation coefficient (ICC) as given in [20] and follow the implementation of [19]. Overall, we can see there there are varying degrees of ICC on both image and patient level. Transformation of the values can help to mitigate the effect of the ICC to some degree (e.g., log10 transformation of "area"), but there is still correlation present after transformation (Fig S3a).

**Simulation for patient and slide effect** To investigate the impact on modelling, we adapted code from [19]. We compare how ICC affects results of linear models, where all observation are treated as independent, compared to linear mixed-effects models where the group-level effects are estimated and accounted for at different levels. Notably, in the [5] dataset, failing to account for the correlation on slide and patient level led to a false discovery rate of up to 20 % for certain metrics. Accounting for correlation using mixed-effects models however, helped to maintain the FDR below the desired 5% level (Fig S3b). In addition, we performed systematic evaluation with varying ICC strengths for three different conditions. Overall, we found that even if ICC increases in one group / condition, the type 1 error increases notably when using linear models while linear mixed effect models accounted for these effects (Fig S3c).

#### Methods comparison

In order to compare the performance of `sosta` with other methods, we performed a systematic comparison based on simulations. We chose simulations since manually annotated datasets often bear some expert subjectivity (c.f., [21]) and simulations allowed us to explore different parameter settings and subsequent performance.

##### Simulations

First, using the `simulateTissueBlobs` implementation in `sosta`, we generated a binary image of size $n \times n$ pixels. It further takes two parameters as inputs `n_blobs` and `blob_sd`. The first parameter determines the number of random seeds and the second the bandwidth of a Gaussian smoothing kernel that connects the random seeds. The result is binary image that represents domain and non-domain regions (Fig S4a). Based on this ground truth binary image, we used the function `createPointPatternTissue` to sample cells within the domain and in non-domain regions. We used a Poisson point process (implemented in `spatstat`; [22]) to sample cells. The parameter `lambda_A` controlled the intensity of the point process in the domain regions and the parameter `noise_A` the intensity of the background cells (or noise) in non-domain regions. The cells were allocated a target mark "A". The function `createPointPatternTissue` also generates cells of other marks ("B" and "C") for which the intensities were kept fixed and ignored in the comparison (Fig S4b). We simulated a range of parameter settings. For each parameter combination, we repeated simulations. Each simulation was assigned a random seed to ensure reproducibility. In total, we simulated 576 images.

From the simulated point patterns, we constructed `SpatialExperiment` [23] and `AnnData` [24] objects that could be passed to the different reconstruction methods.

##### Reconstruction strategies

We evaluated methods that take as input coordinates $(x, y)$ of cells and corresponding labels and output a polygon that reflects the reconstructed anatomical regions / structures. In general, the methods choose one of two reconstruction strategies.

**Alpha hull** The alpha hull algorithm calculates a flexible boundary around a set of points [25], and is the generalization of a convex hull to follow concave structures. The algorithm requires a parameter $\alpha$ that determines how tightly the hull wraps around points. Small $\alpha$ values result in a tighter boundary where larger values produce a more convex shape. If the value is chosen too small, the resulting structure resembles a star, while if the value is too high, the algorithm ignores the potentially concave nature of the structure and constructs a convex hull. `imcRtools`, `SPIAT` and `CellCharter` all use an alpha hull algorithm to reconstruct structures and internally estimate the $\alpha$ parameter.

**Density based approaches** The other strategy is a (point pattern) density approach. The idea is to estimate a two dimensional smooth intensity profile of the cells of interest and then define a cut-off to extract a binary segmentation mask. This approach requires the definition of a smoothing kernel, the kernel bandwidth and the cut-off threshold; `sosta` and `GRIDGENE` employ a density based approach. While `sosta` offers methods to estimate parameters internally, parameters in `GRIDGENE` have to be manually defined.

##### Evaluated Methods

**sosta** We applied the function `reconstructShapeDensityImage` to the `SpatialExperiment` with default parameter, selection the mark "A" as target for reconstruction. This returned an `sf` polygon object with the reconstructed tissue. All parameters were internally estimated. We used `sosta` version 1.3.2 for methods comparison [8].

**imcRtools** We first constructed a spatial graph with `buildSpatialGraph` using a $k$ nearest neighbourhood definition with $k = 10$. We then identified connected cell patches with `patchDetection`. Polygon boundaries of detected patches were extracted with patchSize and converted to sf objects. We used imcRtools version 1.14.0 for methods comparison [26].

**SPIAT** We used a custom function adapted from identify_bordering_cells to obtain polygons instead of only border cell labels. First the target cells "A" were selected from a SpatialExperiment object, then an alpha hull was computed around the target cells. The alpha parameter was estimated using SPIAT functions. We used a custom function ashape_to_polygon to convert the resulting alpha hull to a sf polygon object. We used SPIAT version 1.10.0 for methods comparison [27].

**CellCharter** The AnnData object was used as input to subset the target mark ("A"), constructed a Delaunay based spatial neighbour graph with gr.spatial_neighbors from squidpy, identified connected components with gr.connected_components, and computed outer boundaries with tl.boundaries. Polygon and multipolygon outputs were converted to WKT strings in Python and then imported into R as sf objects. We used CellCharter version 0.3.7 for methods comparison [28].

**GRIDGENE** For GRIDGENE, we created Python helper functions to convert the point patterns into image like arrays. Point pattern coordinates were rounded to integer pixel locations, and cell marks were encoded into channel specific arrays. We then applied ConvolutionContours using a square kernel. Contours for the target mark "A" were thresholded by local density and filtered by minimum area. Resulting contours were converted to polygons with shapely converted to WKT strings in Python and then imported into R as sf objects. GRIDGENE requires to manually select parameters for the kernel size, the density threshold and the minimal area. We evaluated GRIDGENE over a parameter grid of 120 combinations. We used GRIDGENE version 0.1.0 (commit id eb62b96) for methods comparison [29].

##### Comparison

To compare all methods, we rasterized the resulting polygons into a binary matrix with the same dimension as the ground truth binary image. We then evaluated the performance using the Jaccard index (JI), which is defined as the area of overlap divided by the area of the union of two binary images $A, B$

$$J(A, B) = \frac{|A \cap B|}{|A \cup B|}.$$

We calculated the JI for each image-method combination and across a full parameter sweep for GRIDGENE. We then recorded the highest JI for GRIDGENE as GRIDGENE_best. GRIDGENE_all represents the distribution of all results. We compared the distribution of all JIs using violin and boxplots. Missing values were replaced with 0 and the average reconstruction was compared using a linear model. All differences in mean JI between methods were significant at the 5% level when including 0 for missing values (Fig S4c). Fig S4d shows dependence of the number of target cells and reconsturction performance. We also split the results into low background noise (`noise_A` < 0.01) and high background noise (Fig S4e) to evaluate the effects of noise on reconstruction.

#### Reconstruction and quantification of epithelial crypts in colon tissue

We used annotations of epithelial cells ("epi") as determined by the original publication for reconstruction of epithelial structures (Suppl. Fig S1a). Reconstruction was performed using the function `reconstructShapeDensitySPE` (parameters: bndw = 50, thres = $9 \cdot 10^{-5}$, dim = 2500) of `sosta` (version 1.1.1) [8] (Suppl. Fig S1b).

**Structure metrics** To describe the morphology of the epithelial structures, we calculate the following geometric measurements. The "fibre width" is given by $w = \frac{P-\sqrt{P^2-16A}}{4}$, where $P$ is the perimeter of structure and $A$ is its area. The "shape index" is calculated by $s = \frac{P}{\sqrt{A}}$, where $P$ is the perimeter and $A$ the area of the structure. Geometric calculations were conducted using the `sf` package [9]. Association of structure metrics with histological regions, cell types and the pseudo-time was done using the authors' annotations. For association of structure metrics and pseudo-time, we either assigned a structure metric to each cell by finding the intersection of structures and cell centroid coordinates, or calculated the median pseudo-time per structure.

**Epithelial crypts** To recover crypts, we applied the function `reconstructShapeDensitySPE` of `sosta` to all cells that fell below the density threshold `complement = TRUE` (parameters: bndw = 50, thres = $9 \cdot 10^{-5}$, dim = 2500). To identify transition crypts, we assigned all cells within a certain distance to the crypt boundary to a specific crypt and then calculated the fraction of cell types per crypt. We next considered cell-level annotations from [1] reflecting their association to malignancy (reference, TVA or CRC region), and determined different threshold of composition to identify transition crypts. First, we selected crypts containing between 20 and 500 cells, > 10% reference and > 50% TVA or CRC cells. We also selected "fetal" transition crypts (> 10, > 10% reference and > 10% epithelial fetal like cells), and "proliferative" transition crypts (> 10 cells, > 20% reference and > 30% proliferative like cells) Fig S1d-f). Wile identifying a number of *de novo* transition crypts (Fig 2i) we also identified a lot of false positive crypts (Suppl. Fig S1h-i). Transition crypts are characterised by the spatially organised fusion of distinct cell populations, such that different populations are localised to opposite sides of the crypt. By contrast, most falsely identified crypts contained potentially misannotated cells or cells that were randomly distributed.

#### Reconstruction and analysis of anatomically relevant gradients in germinal centres of the human tonsil

Preprocessing of the 10x Visium HD human tonsil dataset followed recommendations of the online book OSTA (`https://bioconductor.org/books/OSTA/`) [10]. We used the package `SpotSweeper` [2] for initial quality control (Suppl. Fig S2a). Library-size normalization, highly-variable gene selection, and dimensionality reduction was performed using packages `scater` [4] and `scrapper` [11]. We applied `Banksy` to perform spatial clustering of the data [3] (parameters: lambda = 0.5, $k$ = 50, resolution = 0.8, Suppl. Fig S2b). We used `scrapper` and `scater` for aggregation of genes in clusters. Annotation of anatomical regions was done using the function `findMarkers` in `scran` [12], followed by the function `SingleR` [13] to match marker genes to the reference dataset [14] (Suppl. Fig S2c).

**Identification of germinal centres** Germinal centres (GCs) were identified by combining the clusters "germinal centre B cells" (GCBCs) and "activated naïve B cells" (NBCs). Reconstructions were performed using `reconstructShapeDensitySPE` of `sosta` (version 1.1.1) (parameters: thres = $3 \cdot 10^{-4}$) (Suppl. Fig S2d). To define anatomically relevant reference axes, we computed centroids of the GCBC cluster (Fig 2j, red squares) and NBC clusters (Fig 2j, violet squares) and connected them with a linear segment (Fig 2j, white line). Reference anchor points were determined as the side where the linear axis intersected the GC outline on the NBC side. A perpendicular line through each anchor was then calculated to establish a normalized radial distance measure within each GC.

**Identification of gene expression gradients** For subsequent analysis, we selected 16 germinal centres with both GCBC and NBC zones (Fig 2k). The normalized distances within each GC were binned into 30 bins and then aggregated into pseudo-bulk profiles using `scrapper`. Mutual information coefficients (MIC) were calculated using `minerva` [15] to determine genes with a (non)-linear association with the spatial position along the GC axes. Genes with a MIC > 0.999 were selected for visualisation in Fig 2l. Heatmaps of distance-binned and pseudo-bulked expression profiles were generated using `pheatmap` [16] and `ComplexHeatmap` [17]. Exemplary light zone (LZ) and dark zone (DZ) markers in Fig 2l were selected based on published datasets [14,18].
